# Costimulation through TLR2 drives polyfunctional CD8^+^ T cell responses

**DOI:** 10.1101/375840

**Authors:** Fiamma Salerno, Julian J. Freen-van Heeren, Aurelie Guislain, Benoit P. Nicolet, Monika C. Wolkers

**Affiliations:** Department of Hematopoiesis, Sanquin Research-Amsterdam UMC Landsteiner Laboratory, Amsterdam, The Netherlands

**Author notes:** **Correspondence to:** Monika Wolkers. Department of Hematopoiesis, Sanquin Research-Amsterdam UMC Landsteiner Laboratory, Plesmanlaan 125, 1066 CX Amsterdam, The Netherlands. These authors contributed equally to this work (Fiamma Salerno, Julian J. Freen-van Heeren).

## Abstract

Optimal T cell activation requires antigen recognition through the T cell receptor (TCR), engagement of costimulatory molecules, and cytokines. T cells can also directly recognize danger signals through the expression of toll-like receptors (TLRs). Whether TLR ligands have the capacity to provide costimulatory signals and enhance antigen-driven T cell activation is not well understood. Here, we show that TLR2 and TLR7 ligands potently lower the antigen threshold for cytokine production in T cells. To investigate how TLR triggering supports cytokine production, we adapted the protocol for flow cytometry-based fluorescence *in situ* hybridization (Flow-FISH) to mouse T cells. The simultaneous detection of cytokine mRNA and protein with single-cell resolution revealed that TLR triggering primarily drives *de novo* mRNA transcription. *Ifng* mRNA stabilization only occurs when the TCR is engaged. TLR2, but not TLR7-mediated costimulation, can enhance mRNA stability at low antigen levels. Importantly, TLR2 costimulation increases the percentage of polyfunctional T cells, a hallmark of potent T cell responses. In conclusion, TLR-mediated costimulation effectively potentiates T cell effector function to suboptimal antigen levels.

## Introduction

CD8^+^ T cell responses are critical to defend our body from insults. They promote the clearance of primary infections, protect us from previously encountered pathogens, and can control and kill tumor cells. To exert their effector function, T cells produce high levels of effector molecules, such as the key pro-inflammatory cytokines interferon-γ (IFN-γ), tumor necrosis factor-α (TNF-α) and interleukin-2 (IL-2). T cells that produce two or more cytokines, also known as ‘polyfunctional’ T cells, identify the most potent effector T cells against infections and malignant cells (1–3). The induction of polyfunctional T cell responses thus remains the ultimate goal of vaccine and T cell therapy.

T cell effector function as defined by the magnitude of cytokine production depends on three signals: 1) triggering of the T cell receptor (TCR), 2) engagement of costimulatory molecules, and 3) availability of pro-inflammatory cytokines (4–6). All three signals are required for optimal priming of naive T cells (7, 8). In addition, the coordinated engagement of two or more signals can potentiate the activity of effector and memory T cells (9–11). The intensity of TCR signaling can vary depending on the amount and affinity of the antigen (12–14) and, upon suboptimal TCR engagement, triggering of costimulatory molecules, such as CD28, significantly decreases the threshold of T cell activation (15–19). In addition, cytokines like type I IFNs, IL-12 and IL-18 can promote IFN-γ production both on their own, and in combination with TCR engagement (20–23). During infection, T cells are also exposed to danger signals, which can be sensed by pattern recognition receptors, such as Toll-like receptors (TLRs). RNA-seq analysis revealed that long-lived human memory T cells express a strong TLR-signature, which is absent in naive T cells (24). In addition, TLR2 expression on human T cells was significantly increased upon TCR activation (24, 25), and TLR2 engagement was suggested to reduce the minimal TCR threshold required for T cell proliferation and memory formation (26). The acquired expression of TLRs on T cells suggests that TLRs may support the effector function of differentiated T cells. Indeed, we and others showed that effector and memory CD8^+^ T cells - but not naive T cells - can directly respond to TLR2 and TLR7 ligands by producing IFN-γ in an antigen-independent manner (27, 28).

At the molecular level, TLR-triggering alone on T cells specifically drives *de novo* transcription of short-lived *Ifng* mRNA, which promotes a short induction of cytokine production (27). Conversely, the engagement of the TCR also engages post-transcriptional mechanisms that increase the stability of cytokine mRNA and/or drives their translation (27, 29). These post-transcriptional regulatory events are required to reach optimal magnitude and kinetics of cytokine production (29), and can be potentiated by costimulatory signals, such as engagement of CD28 or LFA-1 (30, 31). Whether TLR ligands can also augment cytokine production of TCR triggered effector CD8^+^ T cells and, if so, which mechanisms they employ is yet to be determined.

Here, we show that TLR2 and TLR7 ligands lower the antigen threshold required for cytokine production of effector CD8^+^ T cells, yet by employing different mechanisms of regulation. Whereas costimulation through TLR7 only induces *de novo* mRNA transcription, TLR2 signaling synergizes with the TCR to also prolong the half-life of *Ifng* mRNA. The engagement of mRNA stabilization supported by TLR2 ligands correlates with enhanced polyfunctional capacity of T cells. Thus, our data demonstrate that distinct stimuli can differently integrate with the TCR signaling to fine-tune T cell responses. Unravelling the direct contribution of TLR triggering to T cell effector functions might be exploited in the future to rationalize the use of TLR ligands as adjuvants for vaccination strategies and T cell therapies.

## Materials and Methods

### Mice, human PBMCs and cell culture

Specific pathogen free C57BL/6J mice and C57BL/6J.OT-I TCR transgenic mice (OT-I) were housed and bred in filter top cages in the animal department of the Netherlands Cancer Institute (NKI). Animals used in experiments were 8-12 weeks of age. Experiments were approved by the Experimental Animal Committee (DEC) and performed in accordance with institutional, national and European guidelines. Studies with human T cells were performed in accordance with the Declaration of Helsinki (Seventh Revision, 2013). Buffy coats were obtained from healthy donors with written informed consent (Sanquin).

Murine and human T cells, MEC.B7.SigOVA cells, B16-F10 melanoma cells expressing the C-terminal part of ovalbumin (B16-OVA; (32)) and parental B16-F10 melanoma cells were cultured in IMDM (GIBCO-BRL) supplemented with 8% FCS, 2mM L-Glutamine, 20U/mL penicillin G sodium salts, and 20μg/mL streptomycin sulfate. Medium for mouse-derived cells was supplemented with 15μM 2-mercaptoethanol.

Bone marrow (BM) derived macrophages from C57BL/6J mice were generated as previously described (27) and cultured in RPMI 1640 (GE Healthcare) supplemented as above plus 15% L-929 conditioned medium containing recombinant M-CSF.

### T cell isolation and activation

Murine CD8^+^ T cells were purified from C57BL/6J or C57BL/6J.OT-I splenocytes by negative MACS-selection according to manufacturer’s protocol (Miltenyi CD8 isolation kit; 90-99% purity) or by FACS-sorting (BD FACSAria III Cell Sorter; > 99.9% purity; Suppl Fig 1A).

For the generation of antigen-experienced T cells, 1×10^6^ MACS-purified CD8^+^ OT-I T cells were activated for 20h MEC.B7.SigOVA cells as previously described (27). Activated T cells were harvested, washed, and put to rest for 3-15 days in the presence of 10ng/mL recombinant murine IL-7 (rmIL-7; PeproTech). Resting OT-I T cells were stimulated in serum-free IMDM for indicated time points with 0.1nM to 100nM OVA_257–264_ peptide (GenScript), 5μg/mL Pam_3_CysSK_4_ (Pam3), 10μg/mL R848 (both InvivoGen), 100μg/mL Zymosan (Sigma Aldrich, kind gift from M. Nolte), 1ng/mL recombinant murine IL-12 (PeproTech), or a combination thereof. As a control, BM-derived macrophages (BMM) were stimulated in serum-free IMDM for indicated time points with 5μg/mL Pam3, 100μg/mL Zymosan or 2μg/mL lipopolysaccharide (LPS, InvivoGen).

For *ex vivo* experiments, C57BL/6J CD8αβ^+^ T cells were stimulated for 6 hours with 2μg/mL plate bound anti-CD3 alone, (17.A2, Bioceros), or in combination with 1μg/mL soluble anti-CD28 (PV-1, Bioceros), 5μg/mL TLR2 ligand, 10μg/mL TLR7 ligand R848, or a combination thereof.

For studies with human T cells, peripheral blood mononuclear cells (PBMCs) were isolated by Lymphoprep density gradient separation (Stemcell Technologies). CD8^+^ T cells were then purified from PBMCs by MACS-selection according to manufacturer’s protocol (Miltenyi CD8 isolation kit). CD8^+^ T cells were stimulated in serum-free IMDM for 6 hours with 1μg/mL plate-bound anti-CD3 alone (Hit3a, eBioscience), or in combination with 1μg/mL soluble anti-CD28 (CD28.2, eBioscience), 0.1μg/mL Pam3 or 10μg/mL R848.

### B16 melanoma-T cell co-culture

B16-F10 melanoma cells were loaded with indicated amounts of OVA_257-264_ peptide as previously described (29). OT-I T cells were added to pre-seeded tumor cells for 5h at a 6:1 effector:target ratio. When indicated, 0.1, 1 or 5μg/mL Pam3 or 0.1, 1 or 10μg/mL R848 was added at t=0h to the cultures.

### Flow cytometry and intracellular cytokine staining

For flow cytometry analysis and sorting, cells were washed with FACS buffer (phosphate-buffered saline [PBS], containing 1% FCS and 2mM EDTA) and labeled with monoclonal antibodies anti-CD8α (53-6.7), anti-CD8β (H35-17.2), anti-CD4 (clone GK1.5), anti-CD44 (IM-7), anti-L-selectin (CD62L) (MEL-14), anti-CD11b (M1/70), anti-CD11c (N418), anti-F4/80 (BM8), anti-PD-L1 (clone MIH5), anti-IFN-γ (XMG1.2), anti-TNF-α (MP6-XT22), and anti-IL2 (JES6-5H4) (all from eBioscience) for murine cells, or anti-CD8 (SK1), anti-TNF-α (Mab11) (both BD Biosciences), anti-IFN-γ (4.SB3), and anti-IL-2 (MQ1-17H12) (both Biolegend) for human T cells. Near-IR (Life Technologies) was used to exclude dead cells from analysis. For intracellular cytokine staining, cells were cultured in the presence of 1μg/ml brefeldin A (BD Biosciences) as indicated. For detection of degranulation, anti-CD107a (eBio1D43) was added at t=0h to the culture. Upon stimulation, cells were fixed and permeabilized with the Cytofix/Cytoperm kit according to the manufacturer’s protocol (BD Biosciences). The frequency of IFN-γ, TNF-α and IL-2 producing cells was calculated as percentage of Near-IR^neg^ CD8α^+^ OT-I T cells, Near-IR^neg^ CD8αβ^+^ murine T cells or Near-IR^neg^ CD8^+^ human T cells, unless otherwise specified. PD-L1 expression was evaluated on NearIR^neg^ CD4^+^ CD8α^−^ B16-OVA cells. Expression levels were acquired using FACS LSR Fortessa or FACSymphony (BD Biosciences) and data were analyzed using FlowJo software (Tree Star, version 10).

### Single molecule FISH probes

Single-molecule FISH probes of 20 nucleotides for *Ifng*, *Tnfa*, and *Il2* mRNA were designed according to the manufacturer’s guidelines (LGC Biosearch Technologies). Probes with predicted high affinity for secondary target genes (identified by BLASTN) were discarded when the murine gene skyline of the Immunological Genome Project (http://www.immgen.org) indicated gene expression in T cells. This resulted in 33 probes for *Ifng*, 45 probes for *Tnfa*, and 29 probes for *Il2*. Probes were Quasar 670-labeled. Sequences are available upon request. Binding competition assays were performed with identical unlabeled probe sets (Sigma-Aldrich). Quasar 670-labeled probes for human *PHOX2B* were used as control probes.

### Flow-FISH

The Flow-FISH protocol was adapted from (33). Briefly, T cells were activated with indicated stimuli in the presence of 2μM monensin (eBioscience). Cells were stained for extracellular markers, fixed, permeabilized and intracellular cytokine staining was performed with the Cytofix/Cytoperm kit according to the manufacturer’s protocol (BD Biosciences) in 96-well V-bottom plates as previously described. All buffers, antibodies and reagents were freshly supplemented with 4 units recombinant murine RNAse A/B/C inhibitor/ml prior to use (New England BioLabs). Cells were washed twice with 200μl wash buffer (RNAse free water containing 12.5% formamide (Sigma Aldrich), 2X SSC, and 4 units RNAse inhibitor/ml) and transferred to 1.5 ml LoBind Eppendorf tubes (Eppendorf). Cells were incubated for 16h at 37°C + 5% CO_2_ with 15nM FISH probes in 50μl hybridization buffer (RNAse free water containing 10% formamide, 1X SSC, 0.1 g/ml dextran sulfate salts (Sigma Aldrich) and 40 units RNAse inhibitor/ml). Cells were washed once with 1 ml wash buffer prior to acquisition in wash buffer by flow cytometry.

### Quantitative PCR analysis

Total RNA was extracted using Trizol reagent (Invitrogen). cDNA was synthesized using SuperScript III Reverse Transcriptase (Invitrogen) and quantitative Real-Time PCR was performed with SYBR green and a StepOne Plus RT-PCR system (both Applied Biosystems). Reactions were performed in duplicate or triplicate, and C_t_ values were normalized to L32 levels. Primer sequences were previously described (27).

To determine the half-life of cytokine mRNA, T cells were activated for 3h with indicated stimuli, and subsequently treated with 10μg/ml Actinomycin D (Sigma-Aldrich) for indicated time points.

### RNA-sequencing analysis

RNA-seq data of B16-F10 cells left untreated or treated with IFN-γ, and of blood-derived monocytes were retrieved from the Sequence Repository Archive (SRA, https://www.ncbi.nlm.nih.gov/sra) (respectively: Geo: GSE106390, (34) and samples: SRR5483450, SRR5483451, SRR5483452 from Geo: GSE86573, (35)). Fastq files were obtained with fastq-dump (SRA toolkit version 2.5). Reads were mapped and transcripts per million (TPM) were obtained with Salmon ((36), version 0.10.2). TPM values of TLRs were combined into one count table using basic R (version 3.5.1) functions in R-studio (version 1.1.453), and plotted using GraphPad PRISM (version 7).

### Statistical analysis

Results are expressed as mean ± SD. Statistical analysis between groups was performed with GraphPad Prism 7, using paired or unpaired 2-tailed Student *t* test when comparing 2 groups, or 1-way or 2-way ANOVA test with Dunnett’s or Tukey’s multiple comparison when comparing > 2 groups. *P* values < 0.05 were considered statistically significant.

## Results

### TLR2-mediated costimulation lowers the antigen threshold for cytokine production of CD8^+^ T cells

We previously showed that TLR2 ligands can promote antigen-independent production of IFN-γ in CD8^+^ T cells (27). To determine whether TLR2 can also provide costimulatory signals, we isolated and purified spleen-derived CD8^+^ T cells from C57BL/6J mice, which were activated for 6h with αCD3 alone, or in combination with the TLR2 ligand Pam_3_CysSK_4_ (Pam3). We measured the production of the three key cytokines that define effective T cell responses, i.e. IFN-γ, TNF-α, and IL-2 (37–39).

The TLR2 ligand Pam3 potently increased the percentage of IFN-γ producing T cells compared to αCD3 stimulation alone. In fact, Pam3 costimulation resulted in levels of cytokine production similar to levels reached with αCD28 costimulation (Fig 1A). Combining Pam3 and αCD28 induced even higher levels of cytokine production, indicating that these two triggers acted synergistically (Fig 1A). Importantly, Pam3 costimulation was effective in MACS-purified (>90% purity) and in FACS-sorted (>99.9% purity) CD8^+^ T cells (Suppl Fig 1A-C), again demonstrating that the TLR2 ligand acted directly on the T cells (27). Whereas T cells triggered with Pam3 alone only produce IFN-γ (27), in combination with αCD3, or αCD3/αCD28 stimulation Pam3 activation also induced the production of TNF-α and IL-2 in a subset of responding T cells (Fig 1A, Suppl Fig 1B, C). In line with the expression pattern of TLR2 (25, 27), Pam3 increased the cytokine production specifically in memory-like CD44^hi^ T cells, and not of CD44^low^ naive T cells (Suppl Fig 1B, C).

**Figure 1:**
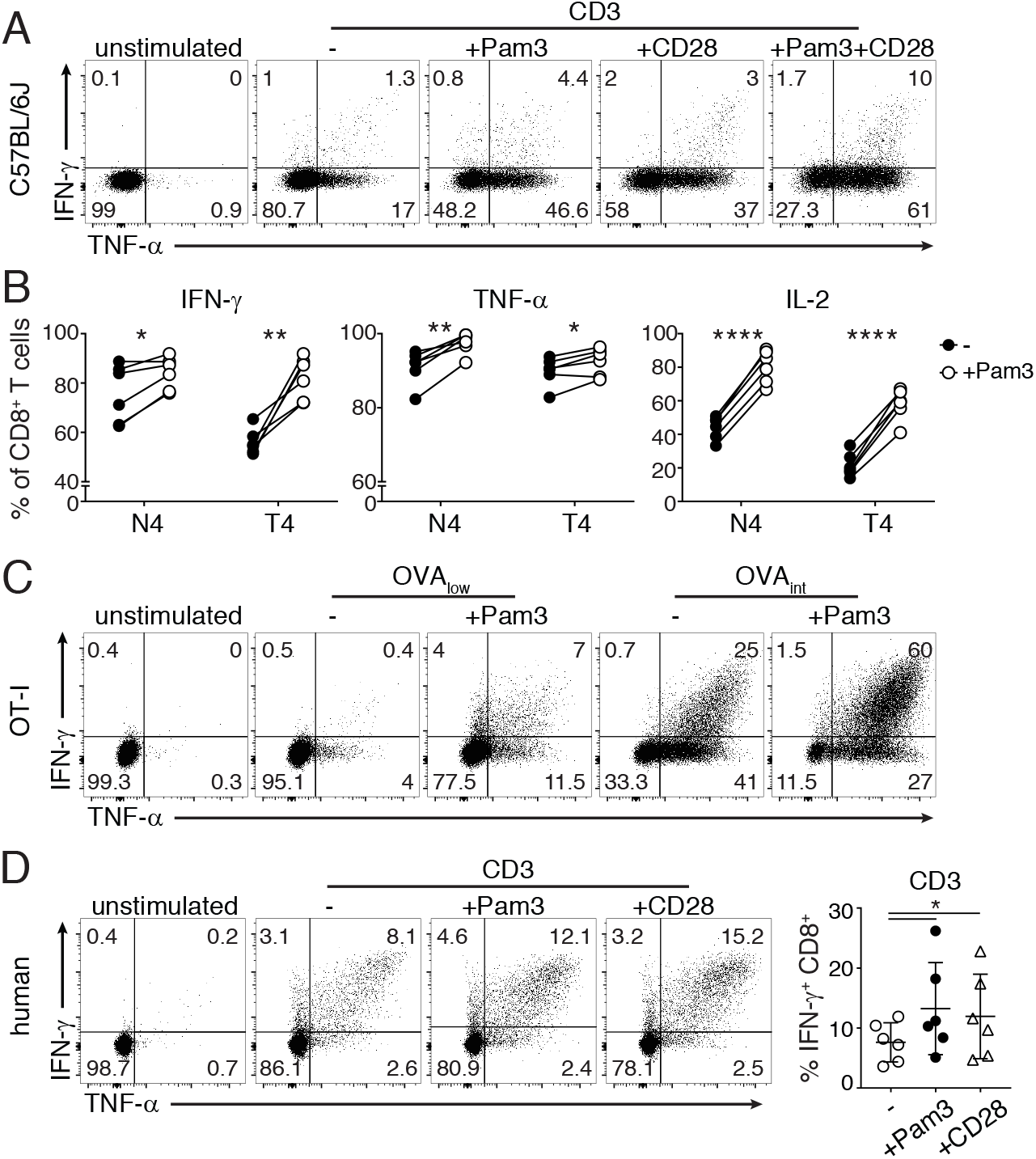
TLR2 acts as a costimulatory molecule and enhances cytokine production of murine and human CD8^+^ T cells. (**A**) CD8^+^ T cells purified from C57BL/6J mice were *ex vivo* stimulated with 2μg/ml αCD3, with or without 5μg/ml TLR2 ligand Pam_3_CysSK_4_ (Pam3), 1μg/ml αCD28, or a combination thereof. Dot plots represent IFN-γ and TNF-α production measured by intracellular cytokine staining. For pooled data see Suppl Fig 1B. (**B**-**C**) *In vitro* activated CD8^+^ OT-I T cells were cultured in rmIL-7 for 3-15 days. T cells were then reactivated with 100nM SIINFEKL (N4) or SIITFEKL (T4) OVA_257-264_ peptide (B), or with 0.1nM (low) or 1nM (int) SIINFEKL OVA_257-264_ peptide (C). When indicated, 5μg/ml Pam3 was added to the cell culture. (B) Compiled data (n=6 mice±SD) of percentage of cytokine producing T cells from two independently performed experiments [Paired Student *t-*test; ^*^p<0.05; ^**^p<0.01; ^****^p<0.0001]. (C) Representative dot plots of 8 mice and 4 independently performed experiments. (**D**) Left: dot plots represent IFN-γ and TNF-α production of human CD8^+^ T cells that were stimulated with 1μg/ml αCD3, with or without 0.1μg/ml TLR2 ligand Pam3 or 1μg/ml αCD28. Right: compiled data (n=6 donors±SD) of percentage of IFN-γ producing T cells from two independently performed experiments [RM-ANOVA with Dunnett’s multiple comparissons collection; ^*^p<0.05]. (A-E) T cells were cultured for 6h in the presence of Brefeldin A. T cells cultured in the absence of stimuli were used as negative control.

To determine the effect of TLR2-mediated costimulation in combination with antigen-specific activation, we turned to TCR transgenic OT-I T cells that were *in vitro* activated and rested for 3-15 days in the absence of antigen, as previously described (27). CD8^+^ T cell purity in this culture system reaches >99.9% at day 3 of culture ((27); data not shown). We stimulated T cells with soluble OVA_257-264_ peptide, allowing for peptide loading of T cells and immediate presentation to the neighbouring cells, without the need to add antigen-presenting cells. This experimental setup enabled us to specifically analyse the direct effect of TLR ligands on T cells. Pam3 costimulation significantly increased the cytokine production of OT-I T cells activated with 100nM OVA_257-264_ peptide (aa:SIINFEKL; N4; Fig 1B). Interestingly, T cells activated with the low affinity variant (aa:SIITFEKL; T4) in combination with Pam3 produced comparable levels of cytokines as when T cells were activated with the high affinity N4 peptide alone (Fig 1B). In line with these results, when Pam3 was added to T cells activated with a low peptide concentration (0.1nM; OVA_low_), IFN-γ and TNF-α production raised from hardly detectable to significant levels (Fig 1C). In addition, Pam3 substantially boosted the protein production of all three measured cytokines in response to intermediate peptide concentrations (1nM; OVA_int_; Fig 1C and data not shown). Thus, TLR2-mediated costimulation with Pam3 lowers the antigen threshold required for T cell activation.

TLR2 dimerizes with TLR1 or with TLR6 (40). To determine whether both two heterodimers induce costimulatory signals to T cells, we activated T cells with OVA_int_ in combination with either Pam3, which activates TLR1/2 heterodimers (41), or zymosan, which activates TLR2/6 heterodimers (42). In contrast to the costimulation induced by Pam3, stimulation with zymosan alone or in combination with OVA_int_ did not result in increased cytokine production (Suppl Fig 2A, B). Nevertheless, zymosan induced TNF-α production of BM-derived macrophages (BMM) (Suppl Fig 2A), demonstrating its biologic activity. Our data thus demonstrate that TLR2 provides costimulation to T cells only when paired with TLR1, and not with TLR6.

Human T cells also express TLR2, which is upregulated upon TCR triggering (25). To determine whether the costimulatory capacity of TLR2 ligands was conserved between mice and men, we purified CD8^+^ T cells from human PBMCs and stimulated them for 6h with αCD3 alone, or in combination with TLR2 ligand Pam3. Pam3 potently increased the percentage of IFN-γ-producing human CD8^+^ T cells to levels of cytokine production that were comparable to CD28-mediated costimulation (Fig 1D). Interestingly, TNF-α and IL-2 production were not impacted upon Pam3 costimulation (Suppl Fig 1E). Altogether, our findings demonstrate that the TLR2 ligand Pam3 provides costimulatory signals to both murine and human T cells.

### Simultaneous analysis of cytokine mRNA and protein production of murine T cells

We next questioned which signals drive the cytokine production of T cells. At the molecular level, the integration of transcriptional and post-transcriptional regulatory events is required to coordinate the production of cytokines (29, 43–47). To dissect the link between distinct T cell signals and regulation of cytokine production, the analysis of both mRNA and protein levels is thus warranted. Because CD8^+^ T cells respond heterogeneously to activation (3, 29, 33), the analysis of bulk populations can mask differences to various stimuli. This may be of particular importance when measuring the effect of costimulatory signals. We therefore reasoned that measuring mRNA and protein on a single cell level would help to better define T cell responses (33, 48).

We recently established a multi-color flow-cytometry-based fluorescence *in situ* hybridization (Flow-FISH) protocol for human T cells that allowed us to simultaneously detect cytokine mRNA and protein with a single-cell resolution (33). Here, we optimized Flow-FISH for mouse T cells (Fig 2A). Upon T cell activation with 100nM OVA (OVA_hi_), all three cytokine mRNAs and corresponding proteins were detectable (Fig 2A). By making use of an unrelated FISH probe set directed against human *PHOX2B* mRNA we verified the specificity of the staining upon T cell activation (control; Fig 2A). Furthermore, when we spiked in increasing amounts of unlabeled cytokine probes to activated T cells, the mRNA signal decreased in a concentration-dependent manner (Suppl Fig 3A).

**Figure 2:**
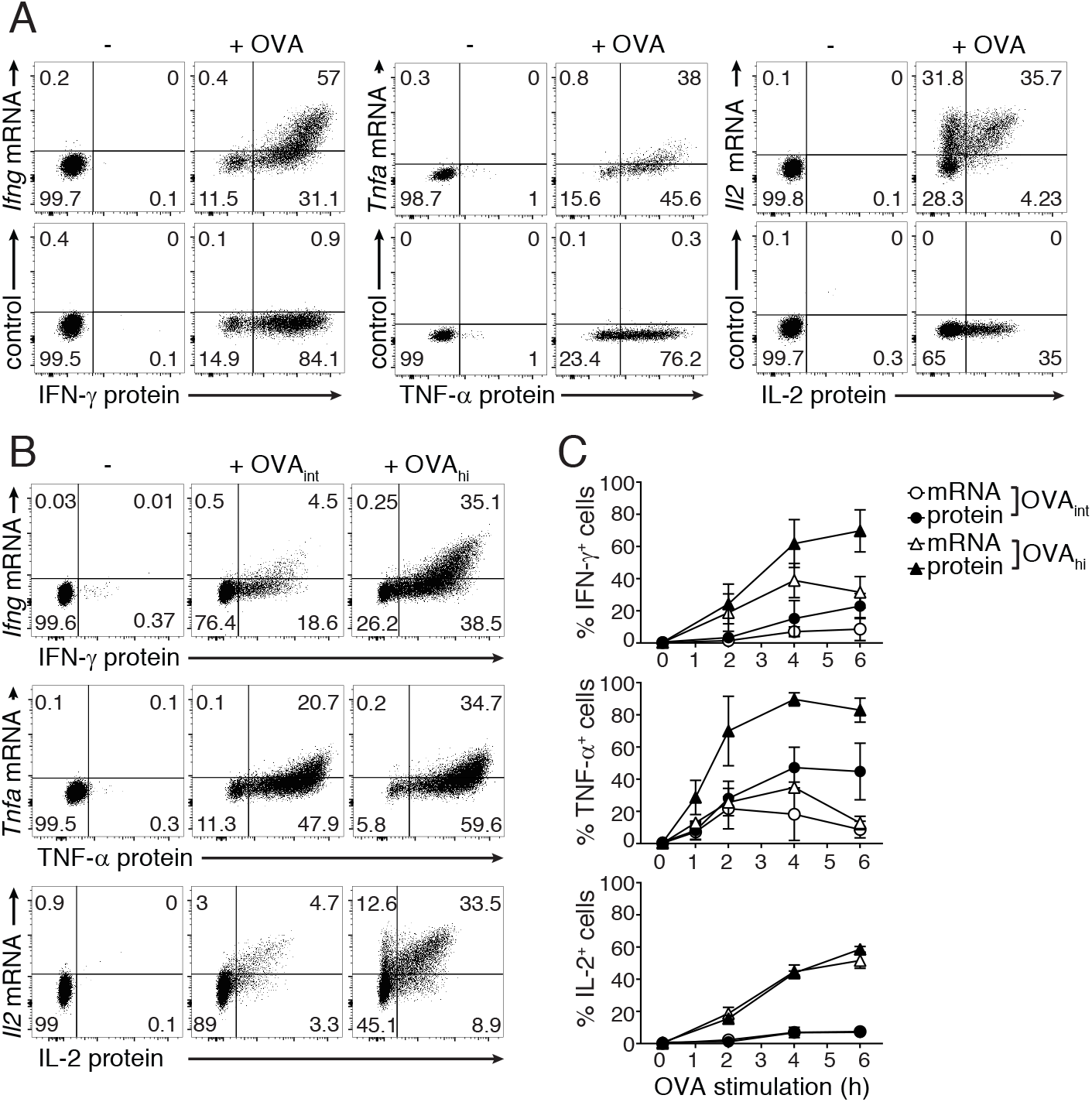
Simultaneous detection of cytokine mRNA and protein by Flow-FISH of murine T cells. (**A**) CD8^+^ OT-I T cells (rested for 5-7 days) were stimulated for 4h with 100nM OVA peptide, and cytokine mRNA and protein expression was analyzed by flow cytometry. T cells were stained with FISH probes specific for *Ifng*, *Tnfa* and *Il2* mRNA (upper panel), or with an unrelated FISH probe set against human *PHOX2B* (lower panel). Dot plots are representative of at least 2 independently performed experiments. (**B-C**) *Ifng*, *Tnfa* and *Il2* mRNA expression and IFN-γ, TNF-α and IL-2 protein production of untreated T cells or T cells stimulated with 1nM (OVA_int_) or 100nM (OVA_hi_) OVA peptide for indicated time points. (B) Dot plots represent cytokine production at 4h of activation. (C) Graphs (mean±SD) show percentage of total mRNA^+^ and total protein^+^ T cells during the entire time course. Data are representative (B) or pooled (C) from 3 independently performed experiments.

We next determined the sensitivity of Flow-FISH. To this end, we turned to our previous findings that cytokine mRNA levels do not always correlate with the actual protein output (29). The production of TNF-α predominately depends on translation of pre-formed mRNA, and only high antigen levels promote *de novo* mRNA transcription. In contrast, the production of IL-2 depends entirely on *de novo* transcription, whereas IFN-γ production depends both on *de novo* transcription and on translation of pre-formed mRNA (29). To determine the capacity of Flow-FISH to detect these differences, we activated T cells with different amounts of antigen and followed their responsiveness over time. As expected, the overall percentage of mRNA^+^ and protein^+^ T cells with Flow-FISH was highest when T cells were stimulated with OVA_hi_ (Fig 2B). Yet, Flow-FISH revealed a cytokine-specific ratio between mRNA^+^ and protein^+^ T cells (Fig 2C). The percentage and the mean fluorescence intensity (MFI) of *Tnfa* mRNA^*+*^cells was comparable whether T cells were stimulated with OVA_int_, or with OVA_hi_ (Fig 2B, C; Suppl Fig 3B). Furthermore, we previously showed that the production of TNF-α protein is driven by increasing translation efficiency, and not by changes in its transcriptional rate (29). This was also reflected by the low levels of *Tnfa* mRNA detected by Flow-FISH (Fig 2B, C). In contrast, irrespective of the antigen load, *Il2* mRNA levels always matched with the corresponding protein levels (Fig 2B, C; Suppl Fig 3B). For IFN-γ, the kinetics of mRNA and protein overlapped when T cells were stimulated with OVA_int_, whereas activation with OVA_hi_ resulted in higher levels of protein^+^ than of mRNA^+^ cells (Fig 2C). These findings show that Flow-FISH is an effective method to visualize the relation of cytokine mRNA and protein on a single cell level.

### TLR2 triggering boosts *Ifng* mRNA transcription

Cytokines follow individual production kinetics upon T cell activation with antigen (3, 29, 33). To determine if costimulation through TLR2 triggering altered the magnitude and/or the kinetics of cytokine production, we measured IFN-γ, TNF-α and IL-2 protein by intracellular cytokine staining. To follow cytokine production over time, we only blocked protein secretion with brefeldin A for the last 2h of stimulation. Pam3 significantly enhanced the percentage of IFN-γ, TNF-α and IL-2 producing T cells upon OVA_int_ stimulation, with the most prominent effect on IFN-γ (Fig 3A). Pam3 mediated costimulation did not affect the kinetics of TNF-α and IL-2 production. Interestingly, the percentage of IFN-γ^+^ T cells after 4h of OVA_int_ stimulation together with Pam3 reached similar percentages as when activated with OVA_hi_ (Fig 3A), but with a significant delay of the onset of production at 2h (OVA_int_+Pam3= 7±2% IFN-γ^+^ T cells, compared to OVA_hi_= 30±8% IFN-γ^+^ T cells). Thus, Pam3 primarily boosts the production of IFN-γ, albeit with a delay in the response kinetics.

**Figure 3:**
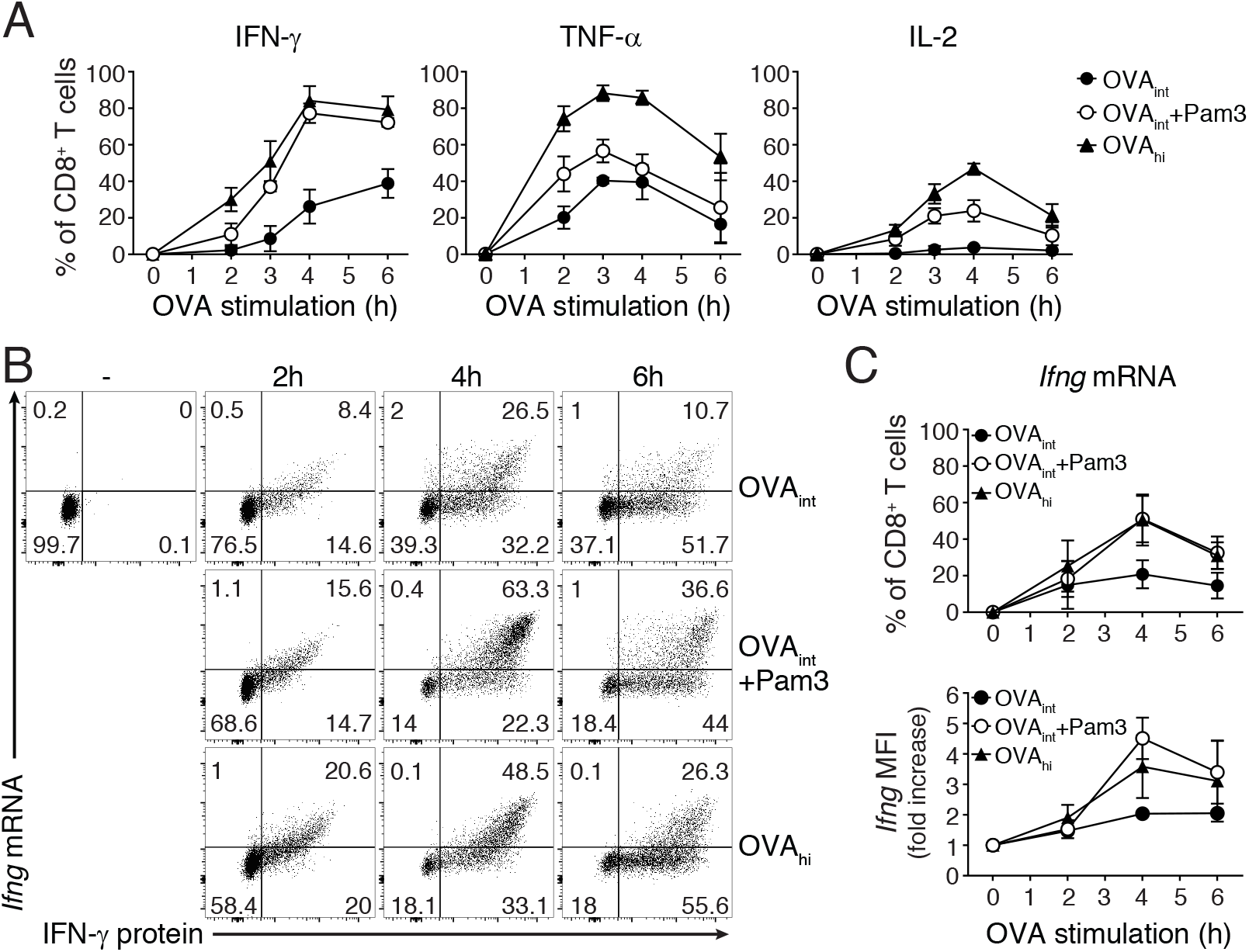
TLR2-mediated costimulation augments *Ifng* mRNA expression levels. (**A**) *In vitro* activated OT-I T cells (rested for 5-7 days) were stimulated with OVA_int_, OVA_int_ plus Pam3 or OVA_hi_ for indicated time points. 1μg/ml brefeldin A was added during the last 2h of activation and intracellular cytokine staining was performed to analyze protein production kinetics. Graphs indicate the total percentage of IFN-γ^+^, TNF-α^+^ and IL-2^+^ T cells (mean±SD) pooled from 3 mice and 3 independently performed experiments. (**B-C**) *Ifng* mRNA expression and IFN-γ protein production was determined by Flow-FISH upon 2, 4, or 6h T cell activation as described above. Unstimulated T cells were used as negative control. (C) Graph indicates the total percentage of *Ifng* mRNA^+^ T cells (top) and fold increase of *Ifng* mRNA MFI upon activation with indicated stimuli compared to unstimulated T cells (time 0; bottom). Data are presented as mean±SD of 3-5 mice and 2-4 independently performed experiments.

We previously found that the TLR2-mediated innate IFN-γ production depends on *de novo* mRNA transcription (27). To determine if similar mechanisms apply for Pam3-mediated costimulation we first measured the cytokine mRNA levels with Flow-FISH. The *Ifng* mRNA levels were not significantly higher in T cells activated for 2h with OVA_int_ with or without Pam3 costimulation (Fig 3B, C). At 4h and 6h of activation, however, the addition of Pam3 significantly boosted the *Ifng* mRNA levels to a similar magnitude as upon stimulation with OVA_hi_ (Fig 3B, C). This was detectable in terms of both the percentage of *Ifng* mRNA^+^ cells and the amount of *Ifng* mRNA produced per cell, as measured by *Ifng* MFI (Fig 3C). Flow-FISH thus showed that TLR-2-mediated costimulation enhances cytokine production by maintaining high mRNA levels.

### TLR2 triggering amplifies the TCR signals to enhance *Ifng* mRNA stability

The magnitude and the duration of IFN-γ production depends on the capacity to stabilize *Ifng* mRNA (29). We determined the mRNA half-life by measuring the *Ifng* mRNA decay from 3h of activation onwards, with RNA polymerase II transcription inhibitor Actinomycin D (ActD) by RT-PCR. Indeed, increased antigen load stabilizes the *Ifng* mRNA (Fig 4A). In line with the limited cytokine production (Fig 1C), activation with OVA_low_ displayed an equally short *Ifng* half-life as unstimulated T cells (t1/2= ~30 min; Fig 4A; (27)). Stimulation with OVA_int_ and OVA_hi_, however, significantly prolonged the half-life of *Ifng* in a dose-dependent manner (OVA_int_: t1/2= ~90 min, OVA_hi_: t1/2 > 2h; Fig 4A).

**Figure 4:**
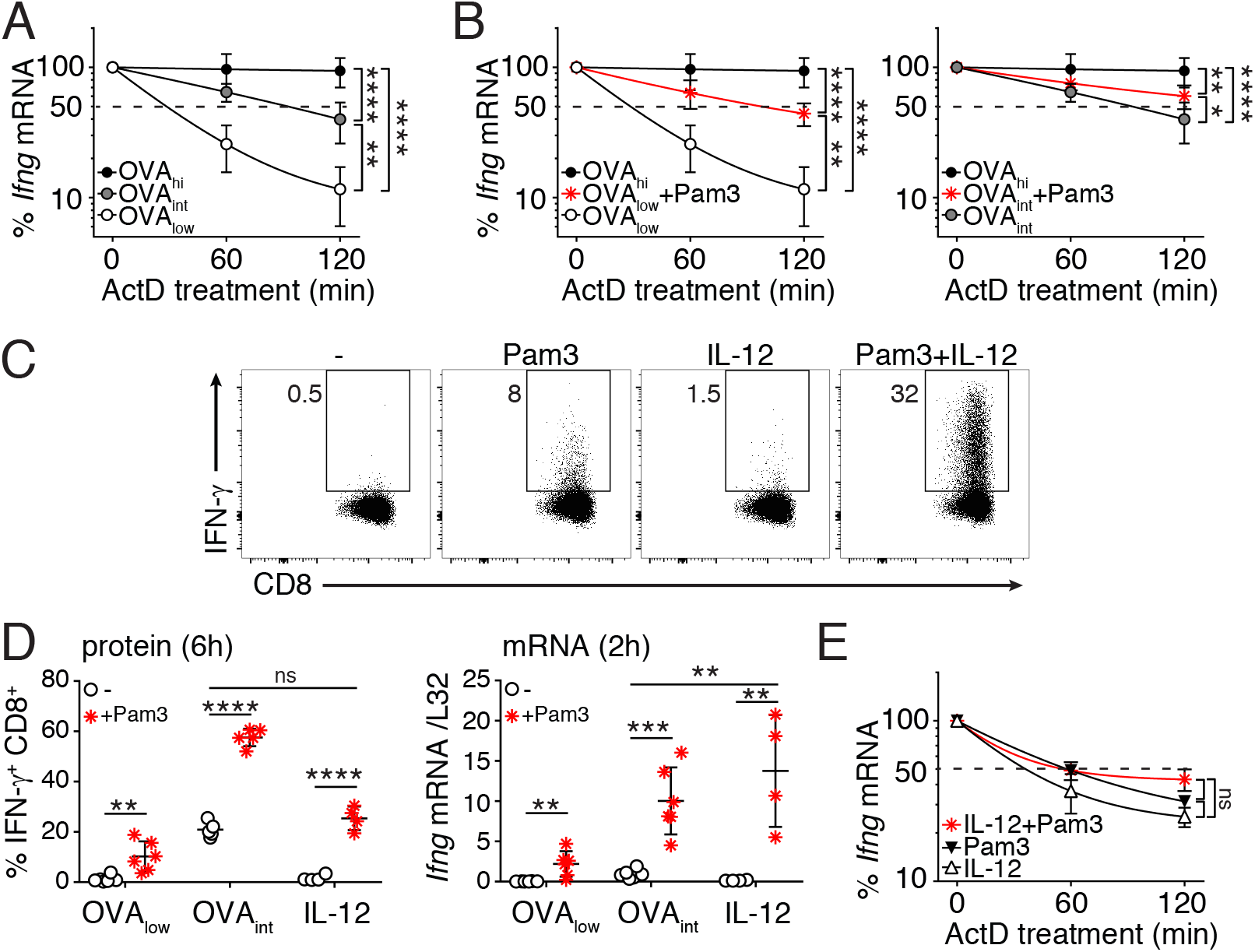
TLR2 triggering enhances TCR-dependent *Ifng* mRNA stability. (**A-B**) *In vitro* activated OT-I T cells (rested for 5-7 days) were stimulated for 3h with OVA_low_, OVA_int_ or OVA_hi_ (A), or with OVA_low_ or OVA_int_ in combination with Pam3 (B), and then incubated with 10μg/ml Actinomycin D (ActD). T cells were harvested at the indicated time points to determine *Ifng* mRNA stability by RT-PCR. Data are presented as mean±SD of 6-8 mice and 4 independently performed experiments [one-way ANOVA with Dunnett’s multiple comparison; ^*^p<0.05; ^**^p<0.01; ^***^p<0.001]. (**C**) Intracellular IFN-γ staining of OT-I T cells stimulated for 6h with 5μg/ml Pam3, 1ng/ml rIL-12, or a combination thereof. Unstimulated T cells were used as negative control. For pooled data see panel D. (**D**) T cells were stimulated with OVA_low_, OVA_int_ or IL-12, with or without the addition of Pam3. Left: graph depicts the percentage of IFN-γ producing T cells after 6h stimulation. Right: *Ifng* mRNA expression was measured by RT-PCR upon 2h T cell stimulation. Data were pooled from 3 independently performed experiments (n=4-5 mice; mean±SD) [Unpaired Student *t*-test; ^**^p<0.005; ^***^p<0.0005; ^****^p<0.0001]. (**E**) *Ifng* mRNA stability of T cells stimulated for 3h as indicated. Data pooled from 4 mice and two independently performed experiments (mean±SD) [one-way ANOVA with Dunnett’s multiple comparison; ns= non-significant].

It was previously shown that costimulation through CD28 and LFA-1 stabilizes cytokine mRNA (30, 31). Stabilization of *Ifng* mRNA was also observed upon Pam3 costimulation. T cell activation with OVA_low_ plus Pam3 increased *Ifng* mRNA half-life to levels similar to OVA_int_ stimulation (t1/2= ~90 min, compare Fig 4A with Fig 4B, left panel). Similarly, OVA_int_ plus Pam3 significantly enhanced *Ifng* mRNA stability (t1/2 > 2h, Fig 4B, right panel).

Because T cell activation with Pam3 alone did not stabilize the *Ifng* mRNA (27), we hypothesized that costimulation itself may not be the determinant factor inducing mRNA stabilization. Rather, we postulated that TCR signaling is required for mRNA stability, and that costimulation can potentiate this process. To study this hypothesis, we activated T cells in a fully antigen-independent manner. When Pam3 stimulation was combined with IL-12, we measured IFN-γ protein levels that were similar to activation with OVA_int_ (Fig 4C, D left panel). Interestingly, at 2h of stimulation Pam3 plus IL-12 induced higher *Ifng* mRNA levels than stimulation with OVA_int_ (Fig 4D right panel). The accumulation of *Ifng* mRNA, however, could not be attributed to stabilization of mRNA (Fig 4E). In conclusion, TLR2 triggering only stabilizes *Ifng* mRNA in the presence of TCR engagement.

### TLR7-mediated costimulation supports the production of IFN-γ without mRNA stabilization

Not only TLR2 triggering, but also TLR7 triggering induces IFN-γ production by T cells in an antigen-independent manner (27). To determine if TLR7 ligands also provide costimulatory signals, we stimulated OT-I T cells with OVA_low_ or OVA_int_ in combination with the TLR7 ligand R848. As TLR2 engagement alike, TLR7 triggering lowered the antigen threshold of T cell activation and significantly increased the production of IFN-γ (Fig 5A). Similar results were found for *ex vivo* FACS-sorted memory-like CD44^hi^ CD8^+^ T cells activated with αCD3 and/or αCD28 and R848 (Suppl Fig 1D). However, in contrast to Pam3, this effect appears to be species-specific, because R848 does not enhance the cytokine production of human CD8^+^ T cells (Suppl Fig 1E).

When turning back to our mouse system, we observed that R848 also supported IL-12-mediated innate production of IFN-γ (Fig 5A, B). The increase in IFN-γ protein^+^ T cells directly correlated with increased *Ifng* mRNA levels (Fig 5B). *Ifng* mRNA consistently displayed a short half-life when T cells were stimulated in an antigen-independent manner (compare Fig 4E and Fig 5C), resulting in similar levels of protein production when Pam3 or R848 were used in combination with IL-12 (Fig 5D). Interestingly, TLR7-mediated costimulation completely failed to prolong *Ifng* mRNA half-life when engaged in combination with the TCR (Fig 5C), and led to lower levels of protein production than TLR2-mediated costimulation (Fig 5D). Altogether, our data show that the benefit of mRNA stabilization requires TCR signaling, can be prolonged by TLR2 ligands and significantly potentiates cytokine production.

**Figure 5:**
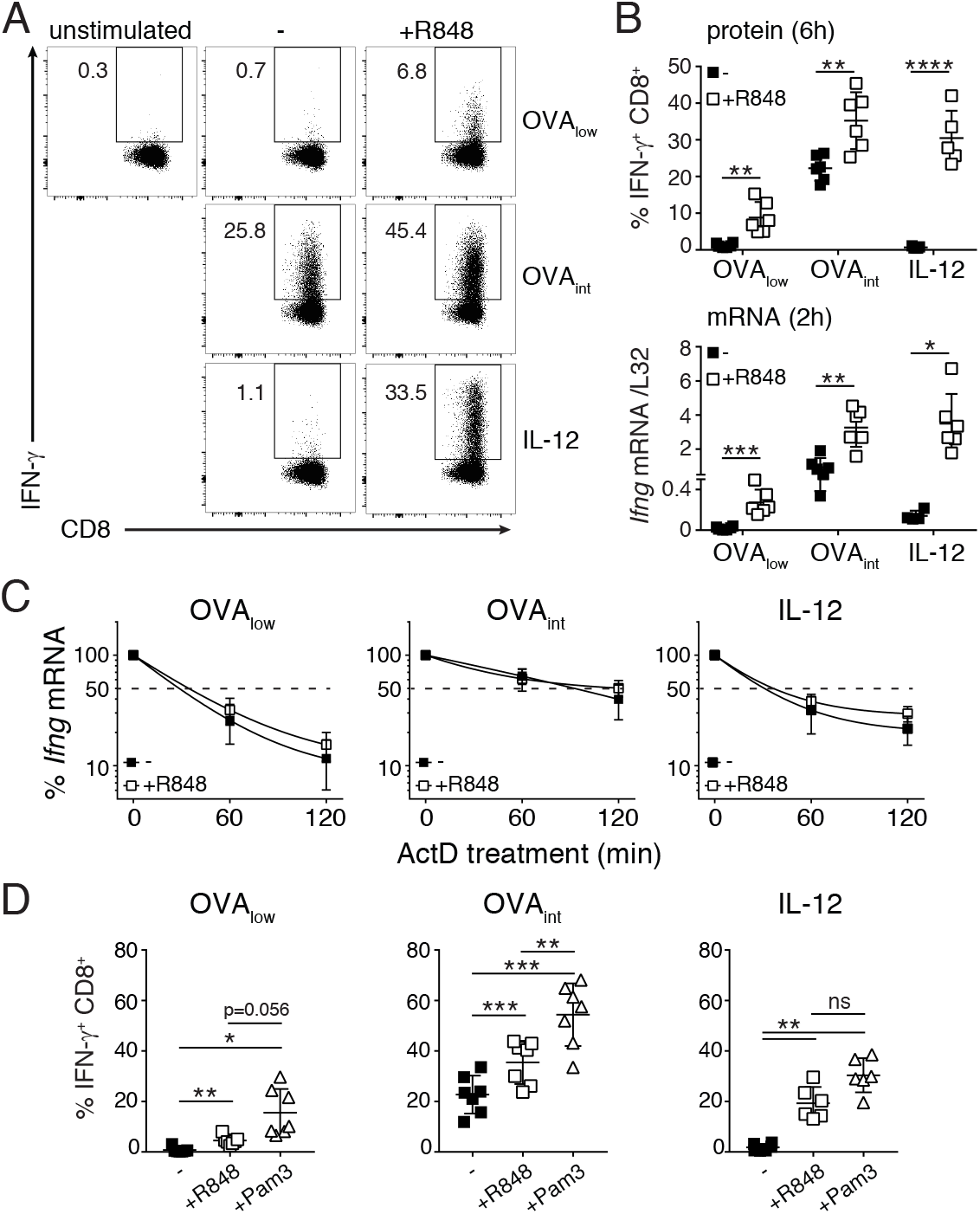
The engagement of TLR7 promotes IFN-γ production of T cells without enhancing *Ifng* mRNA stability. *In vitro* activated OT-I T cells (rested for 5-7 days) were stimulated with OVA_low_, OVA_int_ or IL-12, with or without the addition of 10μg/ml R848. Unstimulated T cells were used as negative control. (**A**) Intracellular IFN-γ staining of T cells stimulated for 6h. For pooled data see panel B. (**B**) Top: IFN-γ protein production. Bottom: *Ifng* mRNA expression of T cells stimulated for 2h, and measured by RT-PCR. Data were pooled from 3 independently performed experiments (n=4-5 mice; mean±SD) [Unpaired Student *t*-test; ^*^p<0.05; ^**^p<0.005; ^***^p<0.0005]. (**C**) *Ifng* mRNA stability was determine upon 3h T cell activation, and addition of ActD for indicated time points. Data were pooled from 4 independently performed experiments (n=6-8 mice; mean±SD). (**D**) IFN-γ protein production after 6h T cell stimulation with indicated stimuli. Graphs show data (mean±SD) pooled from 6 mice and 3 independently performed experiments [one-way ANOVA with Tukey’s multiple comparison; ^*^p<0.05; ^**^p<0.01].

### TLR2-mediated costimulation supports polyfunctional CD8^+^ T cell responses against tumor cells

TLR ligands are often used as adjuvants for immunotherapy to boost innate immune responses and enhance antigen presentation (49–56). We here investigated whether TLR ligands can also directly augment the cytokine profile of T cell responses against tumor cells. We loaded B16-F10 melanoma cells with increasing amounts of OVA_257-264_ peptide, and measured the production of IFN-γ, TNF-α and IL-2 by OT-I T cells after 5h of co-culture with the tumor cells, in the presence or absence of Pam3 or R848. Of note, B16-F10 melanoma cells do not express TLR1, TLR2, TLR6 or TLR7 at detectable levels when compared to blood monocytes (Suppl Fig 4A). We could therefore directly determine the direct effect of TLR ligands on T cell effector function against tumor cells. Whereas TLR costimulation increased the percentage of TNF-α^+^ and IL-2^+^ T cells only at high antigen levels, the production of IFN-γ was also supported at low amounts of antigen (Fig 6A, Suppl Fig 4B). Furthermore, the effects of TLR-mediated costimulation were dose-dependent. Increasing concentrations of TLR ligands potentiated the overall percentage of cytokine-producing T cells in response to B16-OVA cells that constitutively expressed the C-terminal part of ovalbumin (32) (Fig 6B). Interestingly, the expression of the degranulation marker CD107a did not change upon TLR-mediated costimulation (Suppl Fig 4C), indicating that TLR signaling specifically promoted cytokine production.

**Figure 6:**
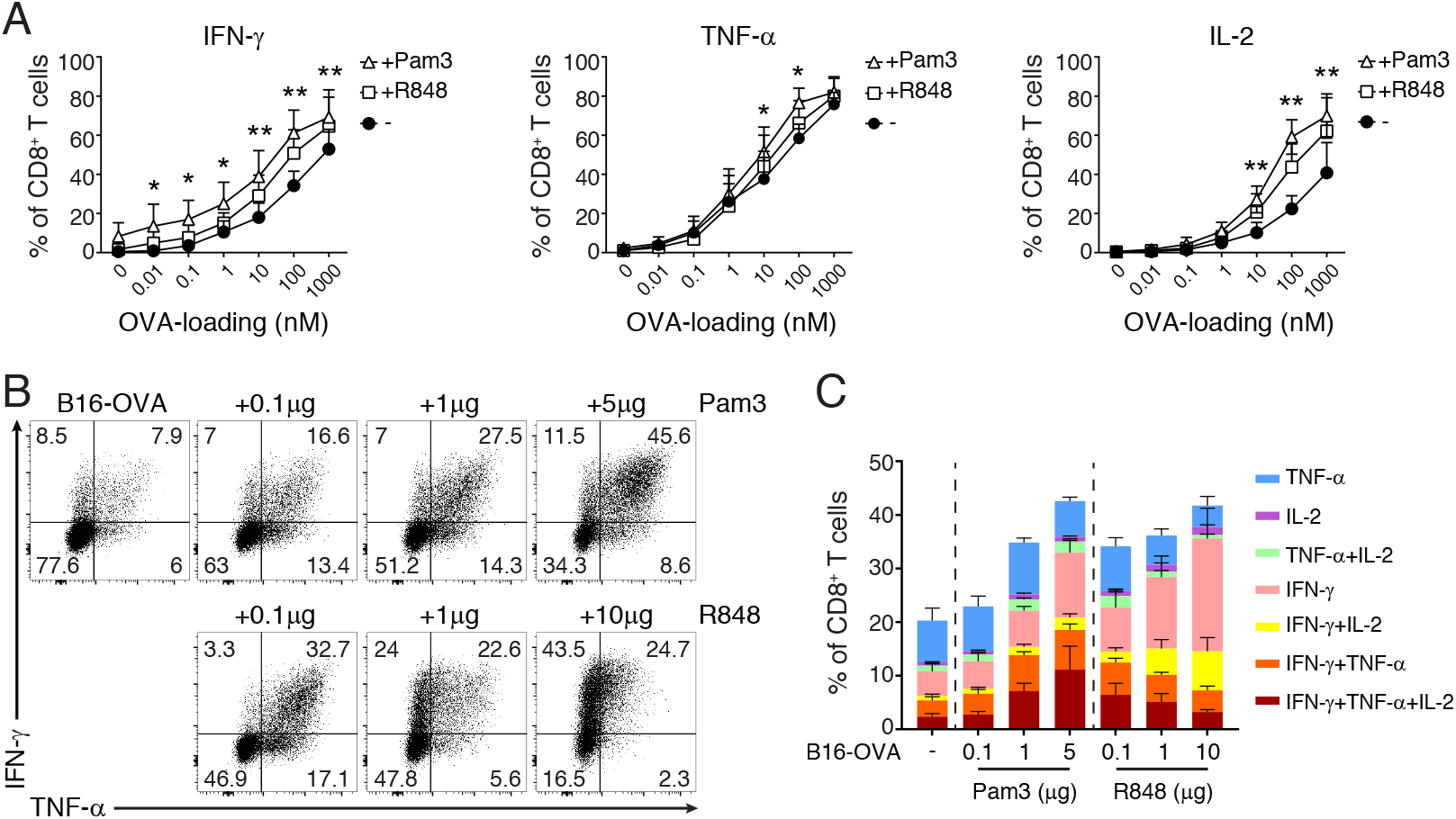
TLR-triggering promotes polyfuntional T cell responses to tumor cells. **(A)** *In vitro* activated OT-I T cells (rested for 5-7 days) were co-cultured at a 6:1 effector:target ratio with B16F10 melanoma cells loaded with indicated amounts of OVA_257-264_ peptide for 6h. When specified, 5μg/ml Pam3 or 10μg/ml R848 were added to the culture system. Graphs represent IFN-γ, TNF-α and IL-2 production of OT-I T cells measured by intracellular cytokine staining. Data is representative of 3 independently performed experiments (n=6 mice; mean ± SD) [two-way ANOVA with Dunnett’s multiple comparison; ^*^p<0.05; ^**^p<0.01]. For representative dot plots, see Suppl Fig 4B. (**B-C**) OT-I T cells were co-cultured with B16 melanoma cells constitutively expressing the C-terminal part of ovalbumin (B16-OVA), as described above. Increasing amounts of Pam3 or R848 were added as indicated. (B) Dot plots represent IFN-γ and TNF-α production measured by intracellular cytokine staining at 5h of activation. (C) Graphs depict the cytokine profile analysis of T cells activated as in B. Data are representative (B) or pooled (C) from 6 mice and three independently performed experiments (mean ± SD).

Potent effector T cells produce two or more cytokines (1–3). This polyfunctional profile correlates with a higher efficacy of vaccines and T cell therapies (38, 57). Intriguingly, even though Pam3 and R848 both supported the cytokine production of T cells in response to tumor cells, the polyfunctional profile of effector T cells was only enhanced by TLR2 ligand Pam3 (Fig 6C). Higher doses of R848 skewed the immune response towards single IFN-γ-producing T cells (Fig 6C). In line with the absence of TLR expression (Suppl Fig 4A), TLR ligands alone did not affect cell viability of tumor cells, or T-cell mediated induction of PD-L1 and MHC-II expression (Suppl Fig 4D, E and data not shown).

In conclusion, TLR2 and TLR7 ligands provide costimulatory signals to CD8^+^ T cells, but they do so through different mechanisms. Importantly, only TLR2 triggering promotes polyfunctional CD8^+^ T cell responses.

## Discussion

CD8^+^ T cells receive different stimuli upon infection that shape their effector functions. This allows T cells to appropriately respond to specific pathogens. Here, we show that TLR2 and TLR7 provide costimulatory signals to amplify the magnitude of cytokine production in murine and in human CD8^+^ T cells. TLR ligands lower the antigen threshold required for T cell activation independently of antigen avidity and/or affinity for the TCR. Whereas antigen-independent engagement of TLRs only induces the production of IFN-γ(27), in combination with TCR triggering, TLR-mediated costimulation also supports the production of TNF-α and IL-2.

In agreement with our previous findings (27), TLR signaling significantly promotes the transcription of *Ifng* mRNA. This effect is further enhanced when TLR ligands synergize with suboptimal antigen levels or with antigen-independent stimuli, such as IL-12. The exact mode of action of TLR-mediated support of TCR and IL-12 signaling is not yet known, and several signal-transduction pathways can be involved. We and others previously showed that TLR2-dependent transcription of *Ifng* mRNA in T cells engages PI3K and Akt signaling (27, 58). In dendritic cells and macrophages, TLR2 and TLR7 engagement also activates nuclear factor kappa B (NF-κB) (59, 60) and activator protein 1 (AP-1) (61, 62), two crucial transcription factors for cytokine production (63, 64). Because PI3K/Akt, NF-κB and AP-1 are also activated through the TCR, we hypothesize that TLR-mediated costimulation can further amplify these pathways in T cells, and synergize with the TCR-dependent transcription factor NFAT to maximize *Ifng* mRNA transcription (63). Interestingly, NF-κB and AP-1 can also be activated downstream of the IL-18 receptor and synergize with IL-12-mediated STAT4 activity to enhance antigen-independent *Ifng* gene expression in T cells (65–67). It is therefore tempting to speculate that the engagement of NF-κB and AP-1 represents the main mode of action of TLR ligands to boost *Ifng* mRNA transcription in T cells, irrespective from the presence of the antigen.

*De novo* transcription of cytokine mRNA is however not sufficient to optimally shape the magnitude and kinetics of cytokine production (29). mRNA turnover is another critical parameter (43–46), and the stability of *Ifng* mRNA strongly depends on the quality and the quantity of stimuli that a T cell receives (27, 29). T cell activation through TLRs alone fails to stabilize *Ifng* mRNA, thereby restricting the magnitude of innate-like cytokine production by T cells (27). Similarly, suboptimal antigen levels alone cannot promote the stabilization of *Ifng* mRNA, but this can be induced with additional costimulatory signals. As for classical costimulatory molecules alike (30, 31), we show here that TLR2 signaling synergized with TCR signaling to stabilize *Ifng* mRNA. Interestingly, TLR7 costimulation failed to stabilize *Ifng* mRNA. The triggering of TLR2, but not TLR7, activates the mitogen-activated protein kinases (MAPK) cascade (60), and MAPK signaling can enhance cytokine mRNA stability (68, 69). Whether TLR2 ligands support TCR-mediated stabilization of cytokine mRNA through MAPK signaling remains to be determined. Regardless of the mechanism, costimulation through TLR2 potentiates cytokine production in response to suboptimal antigen loads by enhancing TCR-mediated stabilization of *Ifng* mRNA.

TLR ligands are currently used as adjuvants for vaccines and T cell therapies (49-52, 54-56). In addition to their well-known effect in boosting innate immune responses and enhancing antigen presentation, we reveal that TLR ligands can also directly augment T cell responses. We recently showed that tumor-infiltrating T cells lose their capacity to produce cytokines because they fail to stabilize the cytokine mRNA (70). It is therefore tempting to speculate that TLR2 could promote anti-tumoral T cell responses by providing costimulatory signals to T cells and prolonging cytokine mRNA half-life. This in turn can potentiate the response rate of other immune cell types, as shown for B cells and macrophages (27, 71). TLR ligands may also act on T cell subsets residing at barrier tissues, like tissue resident memory T cells. These cells are routinely exposed to danger signals and non-cognate pathogens, and may thus develop cell-specific regulatory mechanisms. In conclusion, dissecting how TLR signals integrate within T cells to promote optimal cytokine production may be crucial to rationalize the use of TLR ligands as adjuvants and improve the efficacy of vaccines and immunomodulatory therapies.

## Author Contributions

F.S., J.J.F.H., and A.G. performed experiments, B.P. Nicolet provided technical help and performed RNAseq data analysis, and F.S., J.J.F.H., and M.C.W. designed experiments, analyzed data, and wrote the manuscript.

## Acknowledgements

We would like to thank the animal caretakers from the NKI, B. Popovic for isolating murine bone marrow, and E. Mul for cell sorting.

## SUPPLEMENTARY FIGURE LEGENDS

**Supplementary Figure 1: Cytokine production of activated CD8^+^ T cells**

(**A**) Dot plots represent CD8αβ^+^ T cells (left) and CD8β^−^CD11b^+^ and CD8β^−^CD11c^+^ myeloid cells (right) before selection, after MACS-selection, or after FACS-sorting. (**B-D**) MACS-selected (B) or FACS-sorted (C-D) murine CD8αβ^+^ T cells were stimulated for 6h with Pam3 (B-C), or R848 (D), in combination with anti-CD3 alone or αCD3/αCD28. IFN-γ, TNF-α and IL-2 production was measured by intracellular cytokine staining. Graphs depict the percentage (mean±SD) of cytokine producing CD44^hi^CD8αβ^+^ T cells (left) or CD44^low^CD8αβ^+^ T cells (right) (n=2-3 mice) (**E**) Purified human CD8^+^ T cells were stimulated for 6h with αCD3 alone, or in combination with αCD28, Pam3 (left), or R848 (right). Graphs depict the percentage (mean±SD) of cytokine producing CD8^+^ T cells. Data ± SD are pooled from 6 donors and two independently performed experiments.

**Supplementary Figure 2: TLR1/2-, and not TLR1/6-mediated costimulation enhances T cell cytokine production.**

(**A-B**) *In vitro* activated OT-I T-cells (rested for 3 days) and BM-derived macrophages (BMM) were activated for 4 hours with indicated stimuli in the presence of brefeldin A. (A) Representative dot plots of IFN-γ, TNF-α and IL-2 production of OT-I T-cells (top) and of TNF-α production of BMM (bottom). (B) Graphs show the percentage of cytokine producing OT-I T-cells pooled from 3 mice (mean±SD).

**Supplementary Figure 3: Validation of Flow-FISH analysis of murine CD8^+^ T cells**

**(A)** *In vitro* activated OT-I T cells (rested for 5-7 days) cells were stimulated for 4h with 100nM OVA peptide, and cytokine mRNA and protein expression was measured by Flow-FISH. The specificity of *Ifng*, *Tnfa*, and *Il2* FISH probe signal was determined by adding increasing amounts of competing unlabeled probes during the staining procedure. **(B)** Histograms represent mRNA (left) and protein (right) levels of IFN-γ, TNF-α and IL-2 production upon stimulation of T cells with OVA_int_ (top) or OVA_hi_ (bottom) for indicated time points. Graphs are representative for 3 independently performed experiments.

**Supplementary Figure 4: Cytokine production of OT-I T cells upon co-culture with B16 melanoma cells and the effect of TLR-ligands on B16-OVA cells.**

(**A**) Transcripts per million (TPM) of TLR1, TLR2, TLR6 and TLR7 expression of B16-F10 cells left untreated or treated with IFN-γ, and of blood-derived monocytes. RNA-seq data were retrieved from the Sequence Repository Archive (SRA, https://www.ncbi.nlm.nih.gov/sra) (ref: GSE106390, (34) and samples: SRR5483450, SRR5483451, SRR5483452 from GSE86573, (35), respectively). (**B**) Representative dot plots of IFN-γ and TNF-α production of previously *in vitro* activated OT-I T cells (rested for 5-7 days) that were co-cultured for 5h with B16-F10 melanoma cells loaded with 0.1nM (low), 1nM (int) or 100nM (hi) OVA_257-264_ peptide, with or without 5μg/ml Pam3 or 10 μg/ml R848. (**C**) Graph depicts the percentage (mean±SD) of CD107a^+^ OT-I T cells that were co-culture for 5h with B16-OVA cells with or without the presence of increasing amounts of Pam3 or R848 (n=4-8 mice). (**D**) Graph indicates the percentage of live B16-OVA cells, defined as Near-IR^−^ CD4^+^, that were cultured for 5h in the only presence of Pam3 or R848. Data were pooled from 3 independently performed experiments (mean±SD). (**E**) PD-L1 expression was measured on B16-OVA cells that were cultured alone (shaded black histograms) or co-cultured with OT-I T cells (shaded red histograms), in the presence of indicated amounts of Pam3 or R848. Numbers indicate PD-L1 Geo-MFI. Data are representative of two independently performed experiments.

